# Increased levels of circulating neurotoxic metabolites in patients with mild Covid19

**DOI:** 10.1101/2022.06.22.497189

**Authors:** Estibaliz Santiago-Mujika, Kevin Heinrich, Sonia George, Colt D Capan, Cameron Forton, Zachary Madaj, Amanda R Burmeister, Matthew Sims, J Andrew Pospisilik, Patrik Brundin, Stewart F Graham, Lena Brundin

## Abstract

SARS-CoV-2 corona virus causes a multi-faceted and poorly defined clinical and pathological phenotype involving hyperinflammation, cytokine release, and long-term cognitive deficits, with an undefined neuropathological mechanism. Inflammation increases the activity of the kynurenine pathway, which is linked to neurodegenerative and psychiatric disorders. We sought to determine whether the kynurenine pathway is impacted in patients with mild COVID-19, leading to elevated neurotoxic metabolites in blood, and whether such changes are associated with pro-inflammatory cytokines. Serum samples were taken from 150 patients and analyzed by ELISA and ultra-high performance liquid chromatography (UHPLC). The data were analyzed using multiple linear regression models adjusted for age and sex. We found increased levels of kynurenine, quinolinic acid and 3-hydroxykynurenine in serum from patients with mild COVID-19, together with increased levels of IL-6, ICAM-1, VCAM-1 and neopterin. The levels of neurotoxic metabolites were significantly associated with key inflammatory cytokines including IL-6 and TNFα. The COVID-19 risk-factor hypertension was associated with the highest levels of neurotoxic metabolites in plasma. These neuroactive metabolites could be part of the pathological mechanisms underlying cognitive impairment during and post-COVID and should be explored as potential biomarkers for long-COVID symptoms.

## Introduction

COVID-19, caused by SARS-CoV-2, can lead to systemic disease, including pneumonia, acute respiratory distress syndrome, and impaired consciousness (1). COVID-19 also leads to an activation of innate and adaptive immune responses which result in a substantial inflammatory response (2). In a certain percentage of patients, symptoms persist over time with the potential of symptoms lasting months and up to years after initial illness. In these patients, there is a predominance for those effected by the illness to present neuropsychiatric symptoms (3). Recent studies have shown an increased risk for acute and long-term sequalae after COVID-19 in both vaccinated and unvaccinated patients (4). However, there is currently a lack of understanding for the factors that drive the neuropsychiatric symptoms present during acute and long-term COVID-19. Thus, there is an urgent need to identify biomarkers that can indicate the disease progression during long-COVID, as well as contribute to the understanding of its underlying pathology.

The kynurenine pathway is the major route for tryptophan (TRP) metabolism, and it contributes to several fundamental biological processes, all converging in energy metabolism. Infections and inflammatory conditions can alter the activity of the enzymes in this pathway (5). Several neuroactive metabolites of the kynurenine pathway that bind neuronal receptors have an underlying role in both neurological and psychiatric symptoms (5). As such, quinolinic acid (QUIN) is an agonist of the glutamatergic *N*-methyl-D-aspartate (NMDA) receptor and kynurenic acid (KYNA) is an antagonist of the same receptor (6). In high concentrations, QUIN induces excitotoxic neuronal death by allowing excessive amounts of calcium to enter the cell (6). While KYNA blocks the cholinergic *α*7 nicotinic receptor, it also antagonizes the glycine site of the NMDA-receptor, thus preventing calcium influx (7, 8).

Another kynurenine pathway metabolite, 3-hydroxykynurenine (3-HK) is also neurotoxic and pro-inflammatory (9). 3-HK promotes reactive oxygen species (ROS) generation through several oxidative conversions (10), and 3-HK can accelerate endothelial cell apoptosis (11). The neurotoxic effects of QUIN and 3-HK are additive (5). Entry of QUIN and KYNA into the central nervous system (CNS) is likely partially prohibited by an intact blood brain barrier (BBB). KYN (the metabolite produced from TRP metabolism) and 3-HK, can freely pass the BBB (12). Within the brain, KYN is metabolized to KYNA by astrocytes (5, 13), or to 3-HK and then further to QUIN by microglia and macrophages (5, 14). SARS-CoV-2 infection is known to affect the KYN levels by inducing the production of the pro-inflammatory cytokine, interferon-gamma (IFN-ɣ) that stimulates the rate-limiting enzyme in the kynurenine pathway, indoleamine-2,3-dioxygenase (IDO), thus affecting KYN levels (15).

As previously mentioned, SARS-CoV-2 can lead to a substantial inflammatory response, which affects the levels of different metabolites from the kynurenine pathway. However, there are also other proteins involved in the inflammatory response, such as the intercellular cell adhesion molecule-1 (ICAM-1) and the vascular cell adhesion protein-1 (VCAM-1), in which we were interested. Both ICAM-1 and VCAM-1 are cell surface glycoproteins that govern immune cell migration to sites of inflammation and T-cell-mediated immunity in tissues (16-18). They are expressed in vascular endothelial cells and in response to inflammation, their expression is induced in epithelial and immune cells (19, 20). ICAM-1 and VCAM-1 play a central role in leukocyte trafficking, lymphocyte activation and several immune responses, and the upregulation of ICAM-1 is a signature event during inflammation (21). A recent study showed that monocyte-derived macrophages drive the inflammatory response to SARS-CoV-2 and long-term changes in inflammatory response of monocytes can be detected in convalescent SARS-CoV-2 patients following mild infection (22).Therefore, it is of interest to determine whether the levels of ICAM-1 and VCAM-1 are affected in patients with mild COVID-19, and whether they are linked to kynurenine pathway activation.

In the current study, we analyzed serum samples from 150 patients, 44 of whom tested positive for COVID-19 but exhibited mild disease and were non-hospitalized. We sought to determine whether the production of neuroactive and neurotoxic metabolites along the kynurenine pathway, as well as several proteins involved in inflammation, is altered in patients with mild COVID-19, as this could be associated to the underlying neuropsychiatry symptoms. Kynurenine pathway metabolites are correlated with the severity and predicted negative outcomes of symptoms in COVID-19 patients; and could potentially serve as biomarkers or predictors of neuropsychiatric long-covid symptoms.

## Results

### Demographics of study participants

Serum samples were taken from 150 individuals, 44 were diagnosed with mild COVID-19 (defined for this purpose as positive, but not requiring hospitalization or treatment) and 106 were controls who tested negative for COVID-19. Demographics of study participants are shown in **Table 1**. The average age of SARS-CoV-2 positive individuals was 44.2 ± 13.1 years and controls were 45.6 ± 13.4 years. 47 females (44.3%) were included as controls and 25 females (56.8%) were enrolled with mild COVID-19. Of all participants, 8 controls (7.5%) and 7 COVID +ve (15.9%) were Asian, 5 controls (4.7%) and 0 COVID +ve (0%) were Black/African, 84 controls (79.2%) and 35 COVID +ve (79.5%) were White/Caucasian, 4 controls (3.8%) and 2 COVID +ve (4.5%) were Other, and 5 control (4.7%) and 0 COVID +ve (0%) opted to not answer.

**Table 1.** Demographics of the patients included in the current study.

| **Mild SARS-CoV-2 Demographics** |  |  |
| --- | --- | --- |
|  | **SARS-CoV-2**<br>**Negative (n = 106)** | **SARS-CoV-2**<br>**Positive (n = 44)** |
| **Age**<br>(Mean + SD) | 45.65 + 13.39 | 44.18 + 13.12 |
| **Sex F (%)** | 47 (44.3%) | 25 (56.8%) |
| **Race n (%)** |  |  |
| Asian | 8 (7.5%) | 7 (15.9%) |
| Black/African | 5 (4.7%) | 0 (0%) |
| White/Caucasian | 84 (79.2%) | 35 (79.5%) |
| Other | 4 (3.8%) | 2 (4.5%) |
| No Answer | 5 (4.7%) | 0 (0%) |

SD: Standard Deviation
F: female

### Higher levels of IL-6 present in patients with mild COVID-19

Individuals with mild COVID-19 demonstrated significantly higher levels of IL-6 when compared to negative controls (ANOVA F: 5.260, p= 0.0028**) (**Figure 1a**). TNF-α levels were not altered when both groups were compared (ANOVA F: 2.347, p= 0.075 ns) (**Figure 1b**). No significant differences were observed for the other inflammatory markers including IL-13 (F: 1.847, p= 0.141 ns), IL-8 (F: 1.223, p= 0.304 ns), IFN-ɣ (F: 2.054, p= 0.109 ns, data not shown). All data was corrected for age and sex.

**Figure 1.**
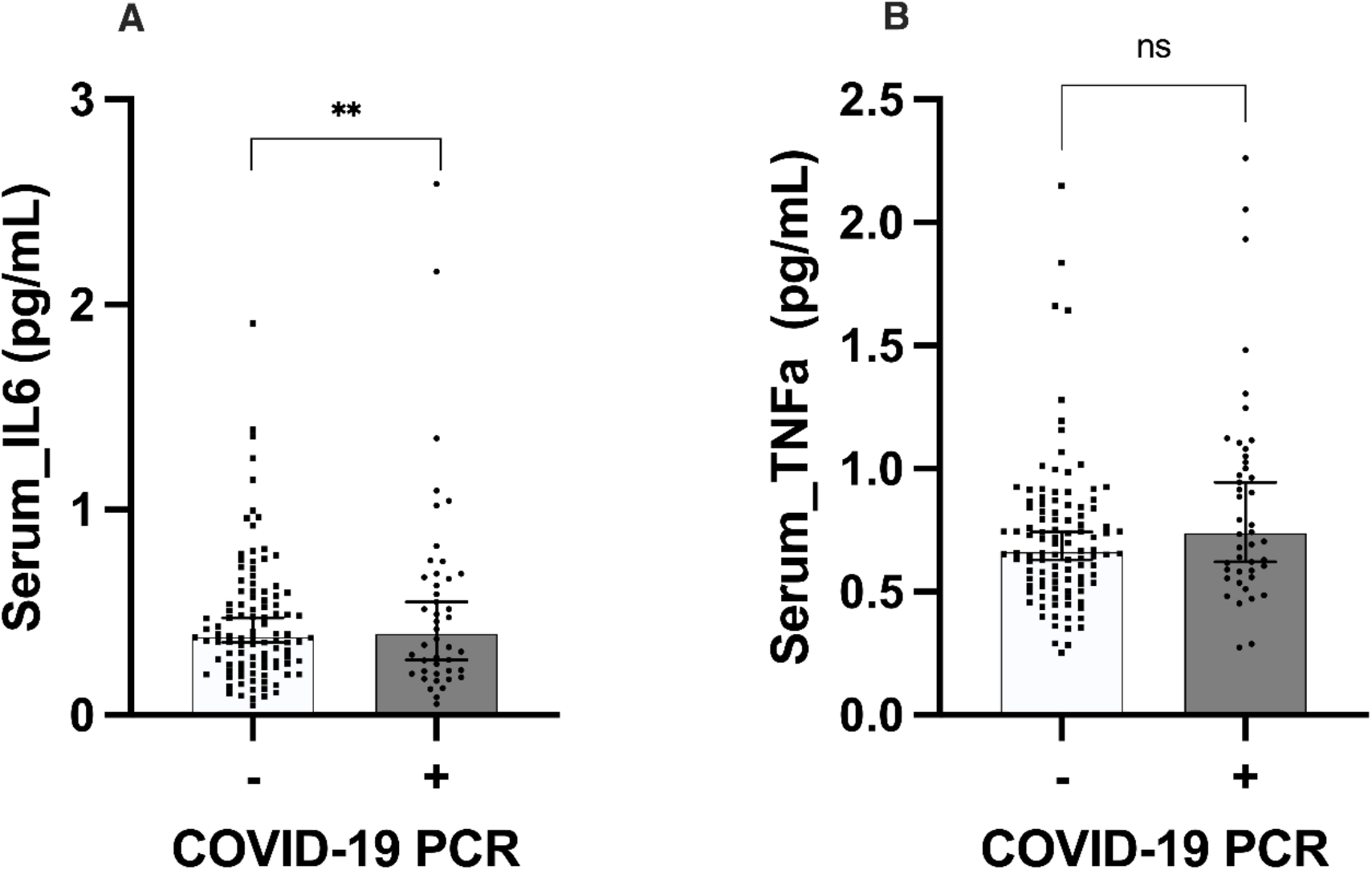
Significantly higher levels IL-6 in patients with mild COVID-19 when compared to controls. **a)** Significantly higher levels of IL-6 were found in patients with COVID-19 when compared to the negative controls (data adjusted for age and sex, ANOVA test F: 5.260, p=0.002 **). **b)** No differences were found in the level of TNF-α between the two groups (data adjusted for age and sex, ANOVA test F: 2.347, p= 0.075 ns). Graphs are represented by median with 95% of confidence interval (CI).

### Higher levels of several proteins related to inflammation present in patients with mild COVID-19

The level of ICAM-1 was increased in patients who tested positive for COVID-19 when compared to the ones that did not (data corrected for sex and age, ANOVA test F: 5.823, p<0.001 ***) (**Figure 2a**). Similarly, patients positive for COVID-19 also showed increased levels of VCAM-1 when compared to the PCR-negative patients (data adjusted for sex and age, ANOVA F: 3.307, p= 0.022 *). The inflammatory related protein, neopterin was also increased in COVID-19 positive patients (data adjusted for age and sex, ANOVA test F: 3.309, p= 0.022 *) (**Figure 2b and 2c)**

**Figure 2.**
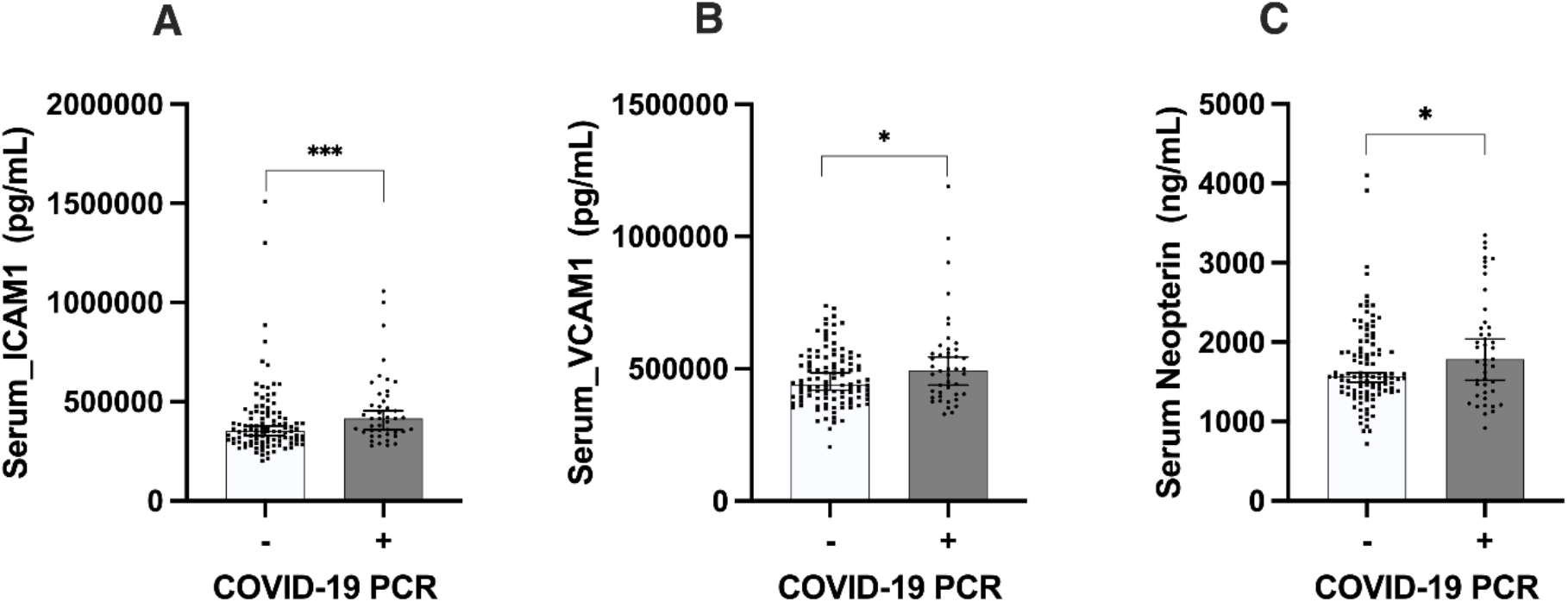
Significantly higher levels of serum ICAM-1, VCAM-1, and neopterin in patients with COVID-19. When the levels of all three proteins were analyzed, patients with COVID-19 presented significantly higher levels of **a)** ICAM-1 (data adjusted for age and sex, ANOVA test F: 5.823, p<0.001 ***), **b)** VCAM-1 (data adjusted for age and sex, ANOVA test F: 3.307, p= 0.022 *), and **c)** neopterin (data adjusted for age and sex, ANOVA test F:3.309, p= 0.022*). Graphs are represented by median with 95% of confidence interval (CI).

### Metabolites of the kynurenine pathway are increased in patients with mild COVID-19

Previous research demonstrated alterations in kynurenine pathway activity following infection (23-25). Therefore, we investigated alterations in kynurenine pathway metabolite levels. Kynurenine (KYN), the first metabolite of the pathway, was significantly increased in patients with mild COVID-19 when compared to patients who tested negative for SARS-CoV-2 (ANOVA test F: 11.195, p= 0<0.001 ***, **Figure 3**). Additionally, 3-HK and QUIN, were increased in patients who tested positive for COVID-19 when compared to patients who did not (ANOVA test F: 3.990, p= 0.009 **; F: 8.492, p<0.001 ***, respectively, **Figure 3**). Further, picolinic acid (PIC) levels in COVID-19 positive patients were significantly decreased when compared to COVID negative patients (ANOVA test F: 4.399, p= 0.005 **). Finally, anthranilic acid (AA) was also significantly increased in patients with mild COVID-19 when compared to the patients who tested negative for the virus (ANOVA test F: 4.024, p= 0.009 **, **Figure 2d**). All data was corrected for age and sex.

**Figure 3.**
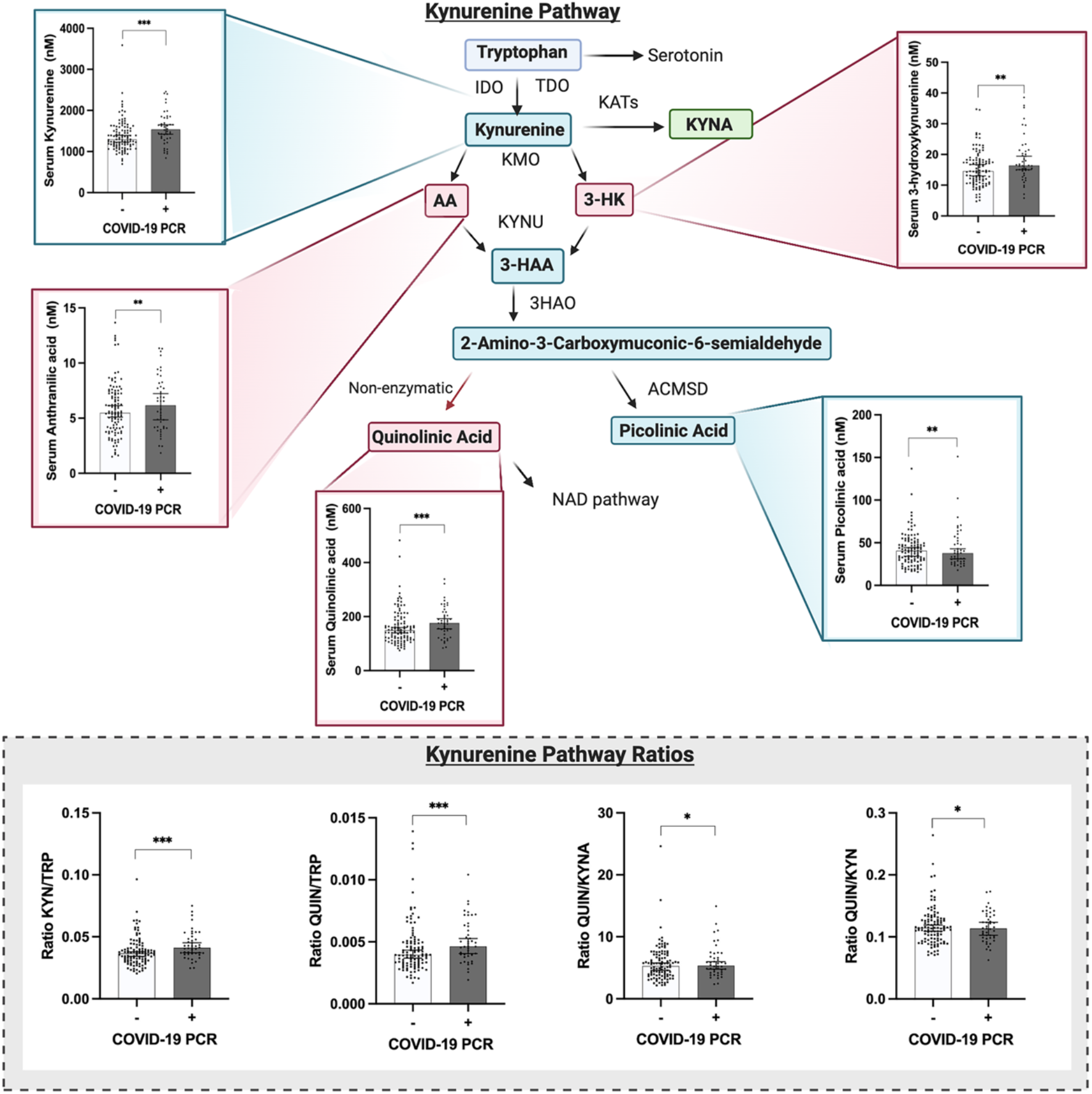
The kynurenine pathway is altered in patients with mild COVID-19, who present increased levels of neurotoxic metabolites. Significantly increased levels of kynurenine (data adjusted for age and sex, ANOVA test F: 11.195, p<0.001 ***), 3-hydroxykynurenine (data adjusted for age and sex, ANOVA test F: 3.390, p=0.009 **), anthranilic acid (data adjusted for age and sex, ANOVA test F: 4.024, p=0.009 **), and quinolinic acid (data adjusted for age and sex, ANOVA test F: 8.492, p<0.001 ***) were found in patients with mild COVID-19 when compared to controls. When the ratio of the metabolites was analyzed, significantly increased levels of KYN/TRP (data adjusted for age and sex, ANOVA test F: 6.377, p<0.001 ***) and QUIN/TRP (data adjusted for age and sex, ANOVA test F: 5.837, p<0.001 ***), as well as QUIN/KYNA (data adjusted for age and sex, ANOVA test F: 2.847, p= 0.040 *) were found in patients with COVID-19. Graphs show the median with 95% of CI. Abbreviations: IDO, Indoleamine 2,3-dioxygenase; TDO, Tryptophan 2,3-dioxygenase; KATs, Kynurenine aminotransferase; KYNA, kynurenic acid; KMO, Kynurenine 3-monooxygenase; AA, anthranilic acid; 3-HK, 3-hydroxykynurenine; KYNU, Kynureninase; 3-HAA, 3-hydroxyanthranilic acid; 3HAO, 3-hydroxyanthranilate oxidase; ACMSD, Aminocarboxymuconate-semialdehyde decarboxylase, and NAD, Nicotinamide adenine dinucleotide.

The ratio of several metabolites was used to further investigate the induction of the kynurenine pathway. The KYN/TRP ratio was quantified to determine the induction of the kynurenine pathway. When both groups were compared, COVID-19 patients showed significantly increased levels of the induction of the kynurenine pathway (ANOVA test F: 6.377, p< 0.001 ***, **Figure 3**). The QUIN/TRP ratio, which has been previously identified as a biomarker for neurological diseases (26), was also analyzed. COVID-19 positive cases showed a statistically significant increase in this ratio when compared to the non-COVID-19 patients (ANOVA test F: 5.837, p<0.001 ***, **Figure 3**). Another neurotoxic ratio that was measured was QUIN/KYNA. In this study, COVID-19 positive patients presented significantly higher ratios when compared to the control group (ANOVA test F: 2.847, p= 0.040 *, **Figure 3**). All data was corrected for age and sex.

### Correlation between neurotoxic metabolites of the kynurenine pathway and pro-inflammatory cytokines

Previous research has shown the relationship between inflammation and the kynurenine pathway activity. Therefore, we subsequently investigated whether the levels of kynurenine metabolites and inflammatory cytokines were correlated. As shown in **Figure 4**, there was a positive correlation between the levels of TNF-α and KYN (Pearson correlation: 0.453; p=0.002**), 3-HK (Pearson correlation: 0.527; p<0.001***) and QUIN (Pearson correlation: 0.482; p<0.001***). Furthermore, a positive correlation between IL-6 and both 3-HK (Pearson correlation: 0.328; p=0.03*) and QUIN (data adjusted for age and sex, Pearson correlation: 0.418; p=0.005**) was also found. All data was corrected for age and sex.

**Figure 4.**
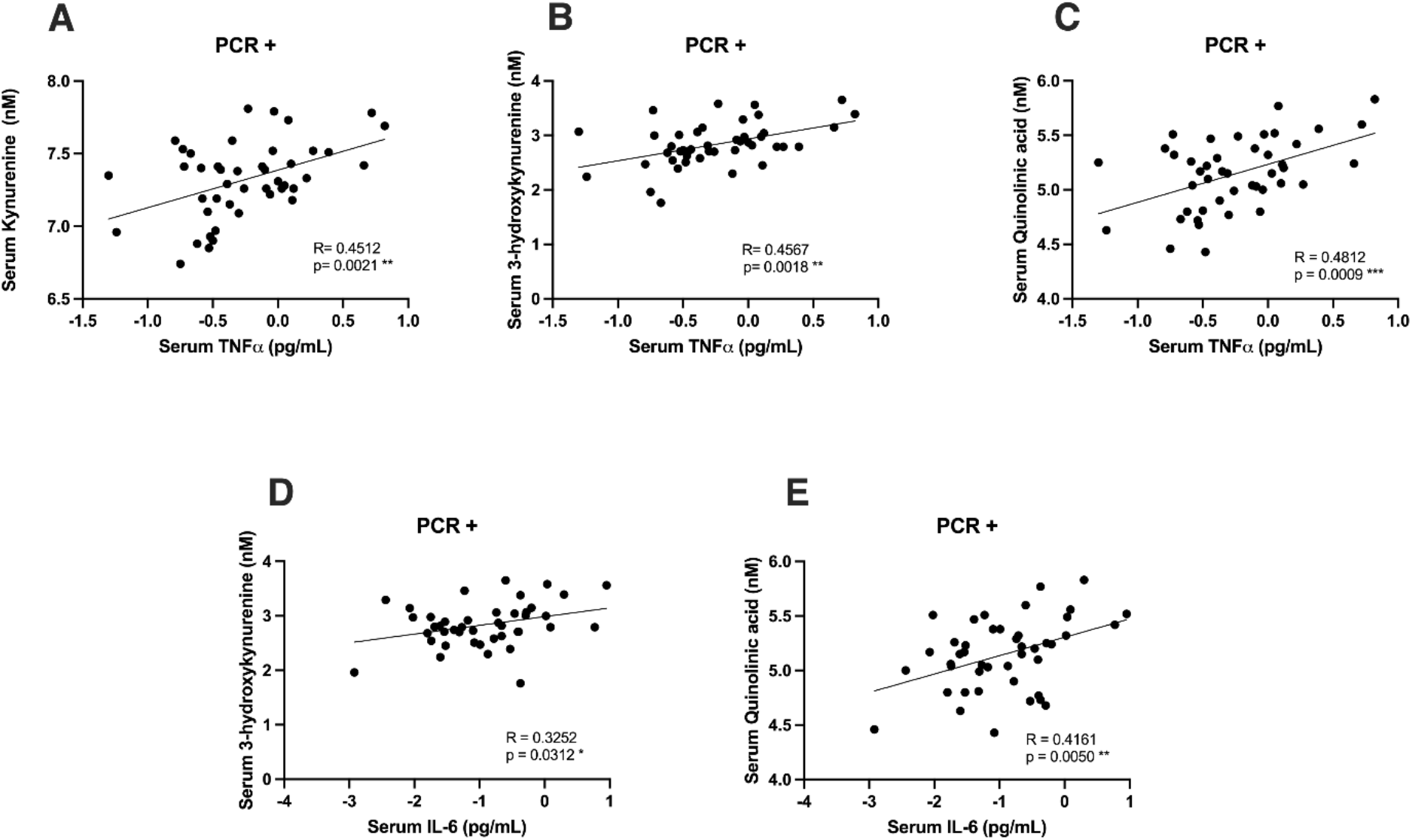
There is correlation between inflammatory cytokines and metabolites of the kynurenine pathway in patients with mild COVID-19. Patients with mild COVID-19 present a positive correlation between TNF-α and kynurenine (data adjusted for age and sex, Pearson R: 0.4512; p=0.0021 **), TNF-α and 3-hydroxykynurenine (data adjusted for age and sex, Pearson R: 0.4567; p=0.0018 **), TNF-α and quinolinic acid (data adjusted for age and sex, Pearson R: 0.4812; p=0.0009 ***), IL-6 and 3-hydroxykynurenine (data adjusted for age and sex, Pearson R: 0.3252, p=0.0312 *) and between IL-6 and quinolinic acid (data adjusted for age and sex, Pearson R: 0.4161, p=0.0050 **).

### Increased levels of neurotoxic metabolites in COVID-19 patients with hypertension

We then analyzed patients who presented with hypertension, a well-known risk factor for severe COVID-19, and compared them with the patients that were not hypertensive in both COVID-19 positive and negative patients. Regression analysis adjusted for age and sex revealed that COVID-19 hypertensive patients presented evidence of higher levels when compared to non-hypertensive COVID-19 patients in the following proteins: IL-2 (fold change 2.66, p=0.003**), IL-6 (fold change 1.45, p-value=0.014*), TNF-α (fold change 1.38, p=0.012*), 3-HK (fold change 1.26, p=0.08), and the QUIN/TRP ratio (fold change 1.14, p = 0.089) (**Figure 5**).

**Figure 5.**
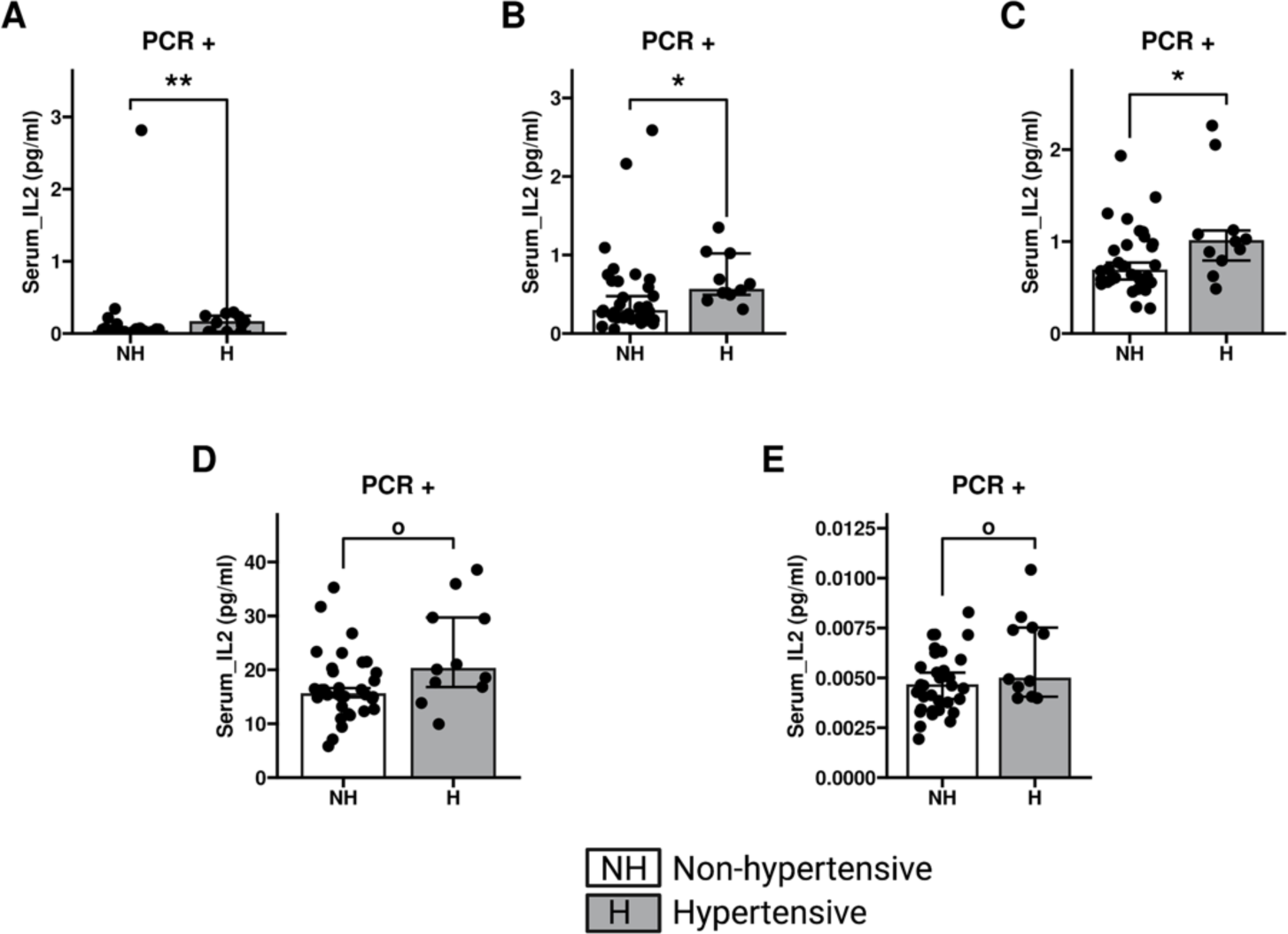
Increased metabolites and ratio in COVID-19 positive hypertensive patients when compared to non-hypertensive COVID-19 patients. Higher levels of IL-2 (fold change 2.66, p=0.003**), IL-6 (fold change 1.45, p-value=0.014*), and TNF-a (fold change 1.38, p=0.012*) in COVID-19 hypertensive patients when compared to COVID-19 non-hypertensive patients. Weak evidence of increased levels in 3-HK (fold change 1.26, p=0.08°), and the QUIN/TRP ratio (fold change 1.14, p = 0.089°). Graphs are represented by median with 95% of confidence interval (CI).

## Discussion

In the current study, we investigated the inflammatory and kynurenine pathway metabolite signatures between mild COVID-19 cases and controls. We found increases in several kynurenine pathway metabolites, such as KYN, QUIN, 3-HK and PIC, together with higher levels of IL-6. Additionally, the increased levels in QUIN/TRP, QUIN/KYN, QUIN/KYNA and KYN/TRP, further support a recent published study by Cihan and colleagues (23). Pro-inflammatory pathway proteins, along with metabolites from the kynurenine pathway were significantly increased in patients that presented with both COVID-19 and hypertension when compared to COVID-19 patients that were not hypertensive. Those proteins included IL-2, IL-6, TNFα, ICAM-1, VCAM-1, 3-HK, QUIN, QUIN:TRP and KYN:TRP.

Kynurenine pathway metabolites are correlated with severity and predicted negative outcomes of symptoms in COVID-19 patients; therefore, it is important to understand the role of kynurenine metabolites in mild COVID-19 patients and long-haulers, in particular those with neuropsychiatric symptoms (15). In this study, we found the levels of ICAM-1 and VCAM-1 to be significantly increased in patients with mild SARS-CoV-2, in particular those with hypertension. Increased levels of ICAM-1 and VCAM-1 in mild and severe cases of COVID-19 infection has already been reported in a small study (27), where it was observed that the severity of COVID-19 disease was associated with increased levels of ICAM-1 and VCAM-1. The main caveat with that study was the small number of patients analyzed. Our new findings support the prior observations and link increases in endothelial cell adhesion molecules to kynurenine pathway metabolites, such as IDO.

IDO is expressed in endothelial cells of vessel walls, and under pathological states, a decrease in IDO leads to an increase in VCAM-1 (28). Therefore, understanding how the kynurenine pathway is involved in inflammatory diseases and how it can alter the levels of ICAM-1 and VCAM-1 will be important to understand, especially in the context of COVID-19.

Lionetto et al., measured the levels of KYN and TRP in the serum of healthy patients, SARS-CoV-2 negative patients, and SARS-CoV-2 positive patients (29). In SARS-CoV-2 positive patients, the KYN/TRP ratio was higher when compared to the negative and the healthy controls (29). We also found an increase in the KYN/TRP ratio in SARS-CoV-2 positive patients when compared to those who tested negative. The KYN/TRP ratio is usually used as an indirect measure of the activity of the IDO enzyme (30); which catalyzes tryptophan. Previous research has demonstrated that IDO is regulated by IFN-ɣ (31, 32). We observed no differences in the levels of serum IFN-ɣ in mild COVID-19 samples. However, IDO activation and gene expression has also been shown to be altered by noncanonical pathways in addition to IFN-ɣ (33). This potentially explains the increased kynurenine pathway activation that we observed in our cohort. The activation of the kynurenine pathway may cause long-lasting inflammation rather than the acute inflammation observed early in COVID-19. Therefore, it will be of interest to follow the kynurenine metabolites as markers in patients with symptoms of long-COVID.

Together with the KYN/TRP ratio, the QUIN/TRP ratio was increased in patients with COVID-19 when compared to controls. In a previous study, Drewes and colleagues suggest that cerebrospinal fluid (CSF) QUIN/TRP ratio could be an early, predictive marker of CNS disease (26). In simian-immunodeficient virus (SIV)-infected macaques, the ratio of QUIN/TRP was significantly increased in cases that led to severe encephalitis (26). Encephalitis is one of the many potential consequences of COVID-19 patients (34); and the QUIN/TRP ratio could be used to determine the outcome of patients. Our data show a significant increase in this ratio in COVID-19 patients, when compared to those that tested negative for the virus. We also found that the ratio between QUIN/TRP was also increased in hypertensive COVID-19 patients, suggesting that patients with hypertension might be prone to a worse outcome.

Cihan and colleagues analyzed kynurenine pathway metabolites and inflammatory cytokines and found a positive correlation between IL-6 and various metabolites from the kynurenine pathway (23). Our study supports a positive correlation between IL-6 and QUIN as well as between IL-6 and 3-HK. Since Cihan’s study analyzed severe and ICU cases of COVID-19 patients, it is possible that the lack of correlation between IL-6 and other kynurenine metabolites in our study is because only mild COVID-19 cases were analyzed. The correlation of IL-6 and kynurenine metabolites could also potentially be used as a biomarker of disease severity.

The correlation between IL-6, IFN-ɣ, and TNF-α and kynurenine metabolites supports the link between inflammation, SARS-CoV-2 and the kynurenine pathway. We found a positive correlation between TNF-α and three metabolites: 3-HK, KYN, and QUIN. TNF-α affects the kynurenine pathway in patients with schizophrenia, as a positive correlation between the levels of TNF-α and KYN has been reported (35). TNF-α may accelerate the formation of KYN from the catabolism of TRP (35). We did not observe differences in TNF-α levels between negative and positive SARS-CoV-2 patients, however, we did find a correlation between TNF-α and KYN levels, similar to what has been reported for schizophrenia (35). In this study, QUIN, 3-HK and KYN were significantly increased in patients with mild COVID-19. Both QUIN and 3-HK are neurotoxic metabolites and were positively correlated with TNF-α and IL-6 in the case of QUIN, and TNF-α in the case of 3-HK, it is possible that the inflammation observed in COVID-19 patients further contributes to the neuronal damage caused by the neurotoxic metabolites.

To summarize, we found an increase in neurotoxic metabolites of the kynurenine pathway in patients with mild COVID-19. Furthermore, the neurotoxic metabolites were correlated with inflammatory markers and vascular injury markers, such as TNF-α, IL-6, VCAM-1 and ICAM-1. We hypothesize that the activation of the neurotoxic branch of the kynurenine pathway might contribute to neurological, cognitive, and psychiatric symptoms experienced in COVID-19 and its aftermath. We suggest that these metabolites should be studied further for their potential as biomarkers of long COVID and as potential contributors to the disease mechanisms underlying long COVID.

## Materials and Methods

### Blood samples

Blood was drawn by venipuncture of the right or left antecubital vein. Blood was allowed to clot at room temperature prior to centrifugation. Serum was aliquoted into cryovials and immediately transferred to -80°C for storage until biological assays.

### Detection of tryptophan, serotonin, and kynurenine metabolites

Serum samples were mixed with extraction solution, briefly vortexed and centrifuged. The supernatant, which contains the metabolites of interest was removed and dried under reduced pressure conditions for ninety minutes in a GeneVac EZ-2 Plus speedvac (SP Scientific, Warminster,PA). Dried down extracts were then resuspended in 0.1% formic acid in Milli-Q water, once resuspended samples were centrifuged through a COSTAR Spin-X 0.22-um filter tube and transferred to an amber vial containing a glass insert.

### Quantification of kynurenine metabolites

Kynurenine pathway metabolites (KYN, KYNA, 3-HK, QUIN, PIC, NTA, NIC, AA), TRP and serotonin were quantified using reverse phase ultra-high-performance liquid chromatography (UHPLC) coupled to a triple quadrupole mass spectrometer (1290 Infinity II LC System, 6470 Triple quadrupole, Agilent Technologies, Santa Clara, CA). Five-microliters of samples were injected onto a Vanguard HSS T3 Pre-column that was connected to an Acquity HSS T3 analytical column. Elution conditions used a combination of Solvent A (0.1% formic acid in LC/MS grade Water) and Solvent B (0.1% formic acid in 90% LC/MS grade acetonitrile) at a flow rate of 0.4mL/min. Agilent Masshunter Quantitative Analysis Software (v9.0, Agilent) was used to analyze and export data.

Intra-assay coefficients of variability (CV) for plasma analytes: TRP 4.1%, KYN 2.5%, KYNA 1.6%, 3-HK 5.0%, QUIN 2.8%, PIC 3.3%, NTA 1.6%, NIC 5.4%, AA 8.4% and 5-HT 2.2%.

Inter-assay CVs: TRP 4.8%, KYN 1.8%, KYNA 1.5%, 3-HK 4.9%, QUIN 1.8%, PIC 1.9%, NTA 1.8%, NIC 6.5%, AA 7.2% and 5-HT 4.9%.

Lower limits of detection (LLOD) were found to be as follows: TRP 36.6 nM, KYN 2.2 nM, KYNA 0.16 nM, 3-HK 0.29 nM, QUIN 4.15 nM, PIC 0.63 nM, NTA 0.98 nM, NIC 0.07 nM, AA 0.98 nM and 5-HT 0.73 nM.

### Quantification of cytokines and alpha-synuclein

Three Meso Scale Discovery (MSD) multiplex kits (Meso Scale Diagnostics LLC, Rockville, MD) were read using a MESO QuickPlex SQ 120 plate reader. Samples were run in duplicate according to the manufacturer’s instructions and sample concentrations were generated through MSD Discovery Workbench 4.0 software. Samples below the average LLOD were denoted as the average LLOD across plates. For α-synuclein, the U-PLEX human α-synuclein kit was used. Samples were diluted 1:8 with sample diluent and generated an inter-plate CV of 3%, an average intra-assay CV of 5.2%, and the LLOD of 0.876 pg/mL.

Using the MSD V-PLEX human vascular injury II kit, we quantified C-reactive protein (CRP), serum amyloid A (SAA), ICAM-1, and VCAM-1. Samples were diluted 1:1000 with manufacturer’s diluent reagent. The inter-plate CV was 10.3% (CRP: 8.3%, SAA: 9.7%, ICAM-1: 15.2%, VCAM-1: 8.1 %), average intra-assay CV 5.6% (CRP 4.2%, SAA 6.5%, ICAM-1 6.6%, and VCAM-1 5.2%), the LLOD were 1.077pg/mL CRP, 12.839 pg/mL SAA, 1.037 pg/mL ICAM-1, and 6.618 pg/mL VCAM-1.

The MSD V-PLEX human proinflammatory I kit was used to determine the level of IFN-ɣ, interleukin (IL)-10, IL-12p70, IL-13, IL-1β, IL-2, IL-4, IL-6, IL-8, and tumor necrosis factor alpha (TNF-α). Samples were diluted 1:1 with sample diluent. Inter-plate CV were calculated (**see Supplementary Table 1**), and the LLOD was IFN-ɣ: 0.087 pg/mL, IL-10: 0.025 pg/mL, IL-12p70: 0.0385 pg/mL, IL-13: 0.209 pg/mL, IL-1β: 0.032 pg/mL, IL-2: 0.024 pg/mL, IL-4: 0.004 pg/mL, IL-6: 0.045 pg/mL, IL-8: 0.017 pg/mL, and TNF-α: 0.049 pg/mL.

### Quantification of Neopterin and S100B

Neopterin ELISA kits were purchased from IBL America (Immuno-Biological Laboratories Inc., Minneapolis, MN). Undiluted serum samples were used following the manufacturer’s protocol. S100B ELISA kits were purchased from Millipore (EMD Millipore, St. Louis, MO) and samples were diluted 1:1. Plates were read using a Tecan Infinite M200 Pro plate reader (Tecan Group Ltd, Männedorf, Switzerland). Sample concentrations were generated using a 4 Parameter Logistic Curve Calculator (AAT Bioquest, “Quest Graph Four Parameter Logistic (4PL) Curve Calculator” 15 Jul. 2021, https://www.aatbio.com/tools/four-parameter-logistic-4pl-curve-regression-online-calculator). The average inter-plate CV for Neopterin was 12.1%, the average intra-assay CV was 2.2%, and the LLOD given by the manufacturer is 0.177 ng/mL. For S100B the average inter-plate CV was 5.0%, the average intra-assay CV was 1.9%, and the LLOD was 2.7 pg/mL as specified by the manufacturer.

### Statistical analysis

Models were adjusted for age and sex in all instances. Robust linear regressions to assess differences between PCR+ individuals with and without hypertension were analyzed via R v 4.1.0 (https://cran.r-project.org/). Correlation analyses and ANOVAs were done with SPSS Statistics (version 28.0.1.0). Graphs were generated using GraphPad Prism version 9.0374 (GraphPad Software, La Jolla, CA). For all tests, statistical significance was considered as p<0.05 and weak evidence as 0.05 < p < 0.1.

### Study approval

This study utilized a subset of samples from the Beaumont Health Large-Scale Automated Serologic Testing for COVID-19 study (36) and was approved by the Institutional Review Board (IRB) at the Beaumont Research Institute Detroit, Michigan, USA (2021-110). The final cohort consisted of 150 individuals (44 Mild COVID-19 cases and 106 controls) randomly selected from the registry. Individuals testing positive for SARS-CoV-2 in a qPCR test at the time of sampling were assigned “Covid19 positive” whereas individuals testing negative in a qPCR test for C SARS-CoV-2 at the time of blood sample were assigned “Covid19 negative” for the purpose of this study.

## Author Contributions

Conceptualization, LB SFG, PB; data curation, ESM, ARB, CDC, CF, KH; formal analysis, ESM, LB, SFG, PB, regression analysis with R, ZM; methodology, LB; visualization, ESM, ARB; writing—original draft, ESM, ARB; writing—review and editing, SG, MXH, JAP, PB, SFG, LB.

## Acknowledgements

We would like to thank the Farmer Family Foundation for their generous contribution, which allowed for this research to be conducted. Additionally, we would like to thank the Van Andel Institute Metabolomics and Bioenergetics Core for providing technical assistance. This project was also supported with Beaumont institutional funds as well as through the Beaumont Health Foundation with philanthropic gifts provided by the following: Sidney & Madeline Forbes, Nathan & Catherine Forbes, Edward C. Levy & the Linda Dresner Foundation, Stephen & Bobbi Polk, Warren & Carol Ann Rose & Family, Elizabeth Rose, Mickey Shapiro & Family, S. Evan & Gwen Weiner and The Hearst Foundations.

## List of abbreviations

3-HK: 3-HydroxyKynurenine
AA: Anthranilic Acid
BBB: Blood Brain Barrier
COVID-19: Coronavirus Disease (2019)
CI: Confidence Interval
CSF: Cerebrospinal Fluid
Glu: Glutamate
ICAM-1: Intercellular Adhesion Molecule 1
ICU: Intensive Care Unit
IDO: Indoleamine 2,3-Dioxygenase
IFN-ɣ: Interferon Gamma
IL: Interleukins
KYN: Kynurenine
KYNA: Kynurenic Acid
PA: Picolinic Acid
PCR: Polymerase Chain Reaction
ROS: Reactive Oxygen Species
S100B: S100-Calcium-Binding Protein B
SAA: Serum Amyloid A
SARS-CoV-2: Severe Acute Respiratory Syndrome Coronavirus 2
SIV: Simian Immunodeficient Virus
TNF-α: Tumor Necrosis Alpha
TRP: Tryptophan
QUIN: Quinolinic acid

## Data availability

Data generated is available upon request.

