## supplemental Table1 for "Increased levels of circulating neurotoxic metabolites in patients with mild Covid19"

**Supplementary information**

**Supplementary Table 1. Inter-plate and intra-assay coefficient of variation of the MSD V-PLEX human proinflammatory I kit**

| | Inter-plate CV (%) | Intra-assay CV (%) |
|---|---|---|
| **IFNγ** | 5 | 5.6 |
| **IL-10** | 5.1 | 9.7 |
| **IL-12p70** | 4.9 | 27.9 |
| **IL-13** | 5.8 | 18.8 |
| **IL-1β** | 2.3 | 32 |
| **IL-2** | 2.9 | 28.4 |
| **IL-4** | 6.1 | 37.3 |
| **IL-6** | 4.1 | 10.3 |
| **IL-8** | 4.5 | 2.6 |
| **TNFα** | 6 | 6.6 |
